# Bisphenol A Disrupts Mitochondrial Functionality Leading to Senescence and Apoptosis in Human Amniotic Mesenchymal Stromal Cells

**DOI:** 10.1101/2024.08.29.610279

**Authors:** Sara Ficai, Andrea Papait, Marta Magatti, Alice Masserdotti, Michael Gasik, Antonietta Rosa Silini, Ornella Parolini

## Abstract

In today’s context, microplastics pollution has become an increasingly pressing issue not only for the environmental fallout but also for the assumed negative effects on human health. It is now well-established that microplastics (>1 mm in size) can enter the human body through ingestion, inhalation, dermal contact and also maternal-fetal transmission. Alarming was the recent findings of microplastics within the human term placenta. Among the degradation by-products of microplastics, Bisphenol A (BPA) has emerged as a hazardous chemical, with potential toxicity at multisystemic level, particularly on the earliest stages of human development. Based on these findings, our study focuses on assessing the impact of BPA on properties and functions of mesenchymal stromal cells isolated from the amniotic membrane (hAMSC) of the human term placenta. The amniotic membrane surrounds the fetus, playing a fundamental protective role toward toxic chemicals and pollutants that the mother may encounter. Our research revealed how exposure to increasing concentrations of BPA compromise mitochondrial functionality in hAMSC, resulting in enhanced production of reactive oxygen species at mitochondrial level (mtROS). This, in turn, leads to the stabilization of p53, which triggers an increased expression of p21 and p27 encoding genes and an imbalance in the genetic expression of Bax and Bcl-2. Additionally, we observed upregulated expression of cytokines and chemokines associated with the senescence-associated secretory phenotype (SASP). The increased oxidative stress, which plays a central role in BPA-mediated toxicity, can trigger the activation of the senescence pathways, or culminate in cell death, due to the overwhelming stress conditions. Therefore, our results provide novel insights into the mechanism of action of BPA and elucidates its impact on the functionality of hAMSC. This underscores the pressing need to reconsider the use of BPA as a plastic additive, mitigating the potential adverse effects on babies.

## Introduction

Plastic is extensively used in modern society and its large-scale production presents a major global challenge (1). Poor waste management practices, with only 9% of plastics being properly recycled worldwide (2), contribute to the widespread dispersion of plastic materials in the ecosystem. These plastics undergo physical and chemical degradation, breaking down into smaller fragments (1), (3). Among them, microplastics (>1 mm in size) (4) have attracted considerable attention due to their persistence and potential harm to ecosystems and human health (5).

Human microplastics exposure primarily occurs through ingestion, followed by inhalation and skin contact. Once inside the body, these microplastics can exhibit toxicity at a multisystemic level (5), and there is a growing concern of their safety issued from the point of view of children’s health (6).

Microplastics toxicity is not solely attributed to the possible physical obstruction within organs but can also be influenced by the presence of additives, incorporated during plastic manufacturing, aimed at enhancing its properties (7). Among these additives, Bisphenol A (BPA) is one of the most investigated, especially for its detrimental effects on the endocrine and reproductive systems (8, 9). Studies have emphasized the pervasive nature of BPA exposure across different body tissues, and BPA was identified in human saliva, blood, breast milk, feces, lungs, liver, and placenta (10). Particularly, the recent discovery of BPA within the placenta has raised significant concerns about its potential impacts on human reproduction and gestation (11). Specifically, human exposure to BPA during pregnancy has been correlated with obstetric complications such as preeclampsia, fetal growth restriction, miscarriage, and premature birth (12), (13), (14), (15).

Despite the growing interest in BPA and its potential health effects, our understanding of how it specifically affects the functionality of the placenta, and consequently fetal development, remains incomplete. Recently, exposure to low doses of BPA during gestation and lactation was shown to be associated with an increased level of pro-inflammatory lymphocytes Th17 in mouse offspring (16). Furthermore, BPA has been linked to pathological changes in the placenta by interfering with its metabolic functions (17).

While few studies have investigated this aspect using trophoblast cells (18), (19), (20), (21), (22) significant knowledge gaps persist regarding its impact on other placental cell types.

Of note, microplastics have been specifically found in the amniotic membrane of the human term placenta (11), and considering the growing evidence that mesenchymal stromal cells (MSC) in placental tissues have a prominent role in generating a functional microenvironment critical to a successful pregnancy (23),(24), it is imperative to understand the effects of BPA exposure on MSC from the amniotic membrane (hAMSC). This will be essential to determine if BPA could negatively impact the ability of hAMSC to maintain a functional environment during pregnancy.

Thus, the aim of our study was to investigate, for the first time, the influence of BPA on hAMSC. Our findings demonstrate that BPA negatively affects hAMSC viability by altering mitochondrial functionality. This alteration is associated with increased production of reactive oxygen species (ROS), which in turn affect the formation of the inflammasome complex and induce senescence. These results offer significant insights into the mechanism of action of BPA on mesenchymal stromal cells, providing new information on its implications for amniotic membrane functionality during fetal development.

## Materials and methods

### Ethics statements

Human term placentae were obtained from healthy women following vaginal delivery or caesarean section at term. Placentae were collected after having obtained informed written consent according to the protocols outlined by the local ethical committee ‘Comitato Etico Provinciale di Brescia’ Italy (number NP 2243, January 19, 2016).

### Isolation and culture of hAMSC

Placentae were processed immediately after collection and hAMSC were isolated as previously described (25). Freshly isolated human amniotic mesenchymal stromal cells (hAMSC) were expanded until passage 1 (p1) and then frozen for future experiments. Expansion was performed by plating cells at a density of 10^4^ cells/cm^2^ in Chang D medium (Irvine Scientific, Santa Ana, CA, USA) supplemented with 2 mM L-glutamine and and 1% P/S, at 37 °C in a 5% CO_2_ incubator. Cells with >98% expression of mesenchymal markers CD13 and CD90, <2% expression of hematopoietic marker CD45, and <2% expression of epithelial marker CD324 were utilized in this study.

### Treatment of hAMSC with BPA

To assess the effects of Bisphenol A (BPA), hAMSC p1 were thawed and seeded at a density of 40 x 10^4^ cells/cm^2^ in Chang D medium supplemented with 2 mM L-glutamine and 1% P/S and maintained in a standard cell culture incubator at 37°C with 5% CO_2_. After cells adhesion to the plastic support, they were exposed to increasing concentrations of BPA for 3, 24 and 48 hours. BPA powder (Sigma-Aldrich #239658) was dissolved in 90% methanol to create an intermediate solution (0.2 mM), which was then diluted in DMEM High Glucose medium (Euroclone, Milan, Italy) supplemented with 20% heat-inactivated FBS, 2 mM L-glutamine, and 1% P/S. hAMSC were exposed to six final concentrations of BPA: 0.05, 0.1, 0.2, 0.3, 0.35, and 0.4 μM BPA. Additionally, hAMSC were exposed to treatment with methanol (used to dissolve BPA) at the highest concentration present in the condition with 0.4 μM BPA. The range of doses was selected based on previous literature investigations to ensure that cells were exposed to environmentally relevant concentrations (26). This selection was extensively addressed in the discussion.

### Evaluation of hAMSC mitochondrial functionality

Mitochondrial activity of hAMSC was assessed using the Thiazolyl Blue Tetrazolium Bromide (MTT) colorimetric assay (Sigma-Aldrich) and the Cell Titer-Glo Luminescent Cell Viability assay (Promega, G7570), following the manufacturer’s protocols. Briefly, cells were plated and treated with BPA as previously described. After 24 hours of exposure, mitochondrial functionality was assessed concurrently using the MTT assay and the Cell Titer-Glo assay. For the MTT assay, cells were incubated for 3 hours with 0.5 mg/mL MTT, followed by overnight dissolution of intracellular MTT-formazan crystals in MTT-lysis solution (50% dimethylformamide in deionized H2O supplemented with 20g SDS). Absorbance was measured at 550 nm using a VictorTM X4 plate reader. ATP production was determined using the Cell Titer-Glo kit, where cells were incubated for 20 minutes with 20 μL of Cell Titer-Glo reagent, then transferred in white support plates, and luminescence was measured using a VictorTM X4 plate reader.

### Evaluation of hAMSC viability, total count and apoptotic rate

To evaluate the impact of BPA on hAMSC viability, cells were harvested 3, 24, and 48 hours after exposure to increasing concentrations of BPA. Subsequently, cells were stained with the eBioscienceTM Fixable Viability Dye eFluorTM 780 (Thermo Fisher Scientific), following the manufacturer’s instructions, to analyze viable (eFluor-negative) and dead (eFluor-positive) cells. Samples were acquired using FACS Symphony A3 BD, and the data were analyzed using FlowJo 10.8 software. The absolute cell count was determined by resuspending all samples in the same volume and acquiring at the same speed for a fixed time within the live cell gate.

Furthermore, to investigate the pro-apoptotic effect of BPA, we analyzed the positivity of hAMSC for FITC-Annexin V-propidium iodide (PI) kit (BD Biosciences) according to the manufacturer’s instructions. After a 15-minute incubation in darkness at room temperature, the samples underwent two washing steps with binding buffer. Samples were acquired using FACS Symphony A3 BD within an hour of staining. Collected data were analyzed using FlowJo version 10.8 software. Cells were categorized into the early apoptotic phase (Annexin V+/PI-), late apoptotic phase (Annexin V+/PI+), and necrotic phase (Annexin V-/PI+).

### Measurement of hAMSC oxidative stress

To assess oxidative stress in hAMSC, cells were thawed and seeded in Chang D medium supplemented with 2 mM L-glutamine and 1% P/S either at a density of 40 x 10^4^ cells/cm^2^ for flow cytometry evaluation or seeded at a density of 25 x 10^4^ cells/cm^2^ in IBIDI chambers (IBIDI, Cat.no #81816) for immunofluorescence. Reactive oxygen species (ROS) were detected using the Mitosox Red fluorogenic dye (Life Technologies #M36008), following the manufacturer’s instructions. Briefly, 3 and 24 hours after BPA exposure, hAMSC were incubated with Mitosox Red (0.7 µM) for 20 minutes at 37°C in the dark. Following incubation, cells were washed twice with PBS and acquired using FACS Symphony A3, and subsequently analysed using FlowJo 10.8 software, or using the MICA platform from Leica. For the acquisition via the MICA platform, we analysed a minimum of 200 cells across three fields for most conditions. However, due to a significant reduction of countable cells at the highest BPA concentrations, we increased the number of fields to achieve a sufficient cell count.

### Gene expression analysis of hAMSC

For gene expression analysis, hAMSC were collected in RLT buffer (Qiagen) after 3 and 24 hours of exposure to increasing concentrations of BPA. Samples were preserved at −80°C until use. Upon thawing, total RNA extraction was performed using the EZ1 RNA Cell Mini Kit (Qiagen, Frederick, MD, USA) in a BioRobot EZ1 Advanced XL Workstation. Subsequently, cDNA synthesis was carried out using the iScript Advanced cDNA Synthesis Kit for RT-qPCR (Biorad, Hercules, California, USA). Then, the cDNA was pre-amplified with SsoAdvanced PreAmp Supermix (Biorad).

Real-time PCR was conducted using the Biorad CFX96 Quantitative Real-Time PCR instrument. The cycling program for real-time PCR was as follows: 30 seconds at 95°C, followed by 40 cycles of 10 seconds at 95°C and 20 seconds at 58°C. Data were analysed using Biorad CFX Maestro 2.2 software (Biorad).

**Table 1.**
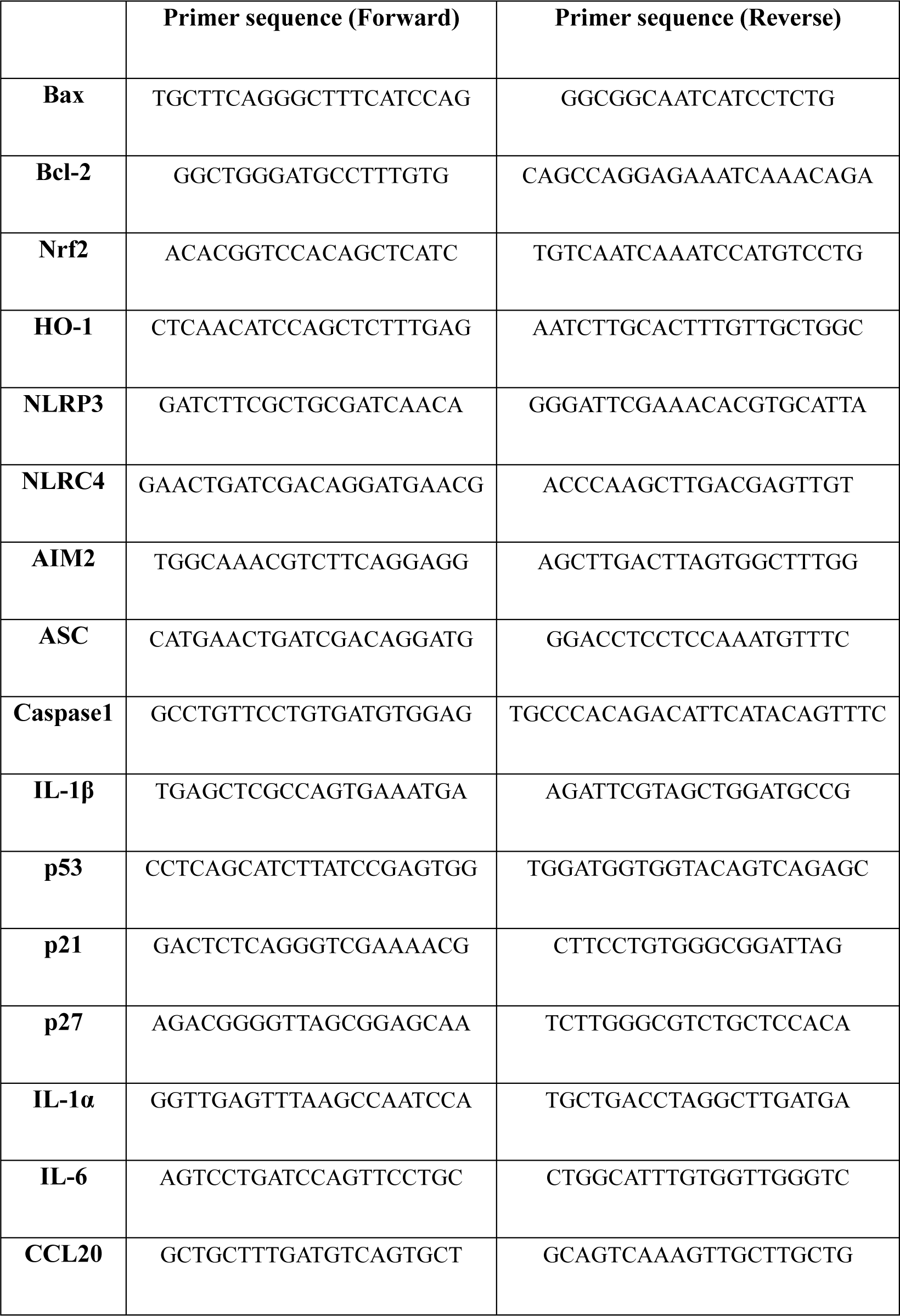
Primer sequences (Sigma Aldrich) of genes analyzed by quantitative real-time PCR.

### Evaluation of hAMSC senescence

To assess hAMSC senescence the Cell Event Senescence Green Detection Kit (Thermo Fisher Scientific, #C10851) was employed following the manufacturer’s instructions. Briefly, hAMSC were seeded onto specific plastic supports for immunofluorescence analysis (IBIDI Cat.no # 81816) at a density of 2.5 x 10^4^ cells/ cm^2^. After exposure to increasing concentrations of BPA for 3 and 24 hours, cells were washed once with PBS 1X and fixed in 4% formalin for 10 minutes. Subsequently, a single wash with PBS 1X was performed, followed by the addition of 100 μL/well of the colorimetric substrate for β-galactosidase (5-bromo-4-chloro-3-indolyl-β-D-galactopyranoside, X-gal) (diluted 1:100 in wash buffer) in the dark for 2 hours at 37°C in normoxia. After the incubation cells were washed once with PBS 1X, and immunofluorescence signals were detected using MICA platform from Leica. A minimum of 100 cells per field for a total of 3 fields, where this was not possible at least 5 fields, were examined to ensure accurate quantification of β-galactosidase activity after treatment.

### Immunofluorescence

hAMSC were seeded onto specific plastic supports for immunofluorescence analysis (IBIDI, Cat.no #81816) at a density of 2.5 x 10^4^ cells/cm^2^. Following 24 hours of exposure to increasing concentrations of BPA, cells were rinsed once with PBS 1X and fixed in 4% formalin for 10 minutes. Subsequently, cells were washed three times with Tris Buffered Saline (TBS) for 5 minutes each.

hAMSC were incubated with p21 primary antibody (Waf1/Cip1/CDKN1A p21, CLONE Santa Cruz Biotechnology), diluted 1:100 in normal goat serum (NGS, Invitrogen, 10000C) and incubated overnight at 4°C in the dark. Following incubation, cells were washed three times in TBS for 5 minutes each, incubated for 30 minutes at 37°C with secondary antibody (VectorLab DyLight 594, ZH1213) at a concentration of 5 µg/mL. hAMSC were then washed again for three times with TBS, and DAPI (0.1 g/mL Cat.no #956) was added for 4 minutes at 4°C in the dark. After one final wash with PBS, images were captured using MICA platform from Leica. A minimum of 200 cells across three fields were analysed for most conditions. However, due to a significant reduction of countable cells at the highest BPA concentrations, we increased the number of fields to ensure an accurate quantification of p21 expression after BPA treatment.

### ELISA assay

Supernatants obtained from hAMSC exposed to increasing concentrations of BPA for 24 hours, with or without an additional treatment of adenosine triphosphate (ATP) for 1.5 hours (1.5 mM), were harvested and stored at −80°C for subsequent analysis. The content of interleukin-1β (IL-1β) was assessed using an enzyme-linked immunosorbent assay (ELISA) kit (BD OptEIA, Cat.no 557953) according to the manufacturer’s instructions. Briefly, 96-well flat-bottom plates (Merck, Cat.no #MSEHNFX40) were coated with 100 μL/well of Coating Antibody and left to incubate for a minimum of 18 hours at 4°C. Afterward, a single wash with Wash Buffer (100 μL/well) was performed, followed by the addition of 200 μL/well of blocking solution (Assay Buffer) for 1 hour at room temperature (RT). Subsequently, samples or standards (50 μL/well) were added to the appropriate wells, along with 50 μL/well of Detection Antibody for IL-1β, followed by a 2-hour incubation at 4°C. Five washes were then conducted, and streptavidin-horseradish peroxidase (HRP) solution was added to each well (100 μL/well) and incubated for 30 minutes at RT. An additional 5 washes were performed with Wash Buffer, and 100 µl/well of 3,3’,5,5’-tetramethylbenzidine (TMB) substrate was added for 30 minutes at RT. Finally, 50 μL/well of Stop Solution was added, and the absorbance was measured at 450 nm using VictorTM X4 (Perkin Elmer).

### Statistical analysis

Data are represented as mean ± standard deviation (SD) in comparison to the control condition (MetOH). Statistical comparisons were performed using a two-way analysis of variance (ANOVA), followed by a Tukey multiple comparison test for post-analysis. The data represent a minimum of three experiments, with the number of replicates (n value) specified in each figure legend. Statistical analysis was conducted using Prism 9 software (GraphPad Software, La Jolla, CA, USA), considering a p-value less than 0.05 as statistically significant.

## Results

### 1. Increasing concentrations of BPA reduce hAMSC viability and trigger an impairment in mitochondrial functionality

We first investigated how increasing concentrations of BPA affect the viability of hAMSC. The experimental control was methanol (MetOH), the solvent employed for dissolving BPA. We observed that 24 hours of exposure with increasing concentrations of BPA caused noticeable morphological changes in hAMSC (Figure 1A). Cells exposed to 0.05 and 0.1 μM BPA exhibited morphology similar to MetOH-treated cells and to cells maintained in the DMEM culture condition. On the other hand, cells exposed to higher concentrations (0.2, 0.3, 0.35, and 0.4 μM BPA) lost the characteristic spindle-like morphology of mesenchymal cells, transitioning to a rounded shape (Figure 1A).

**Figure 1.**
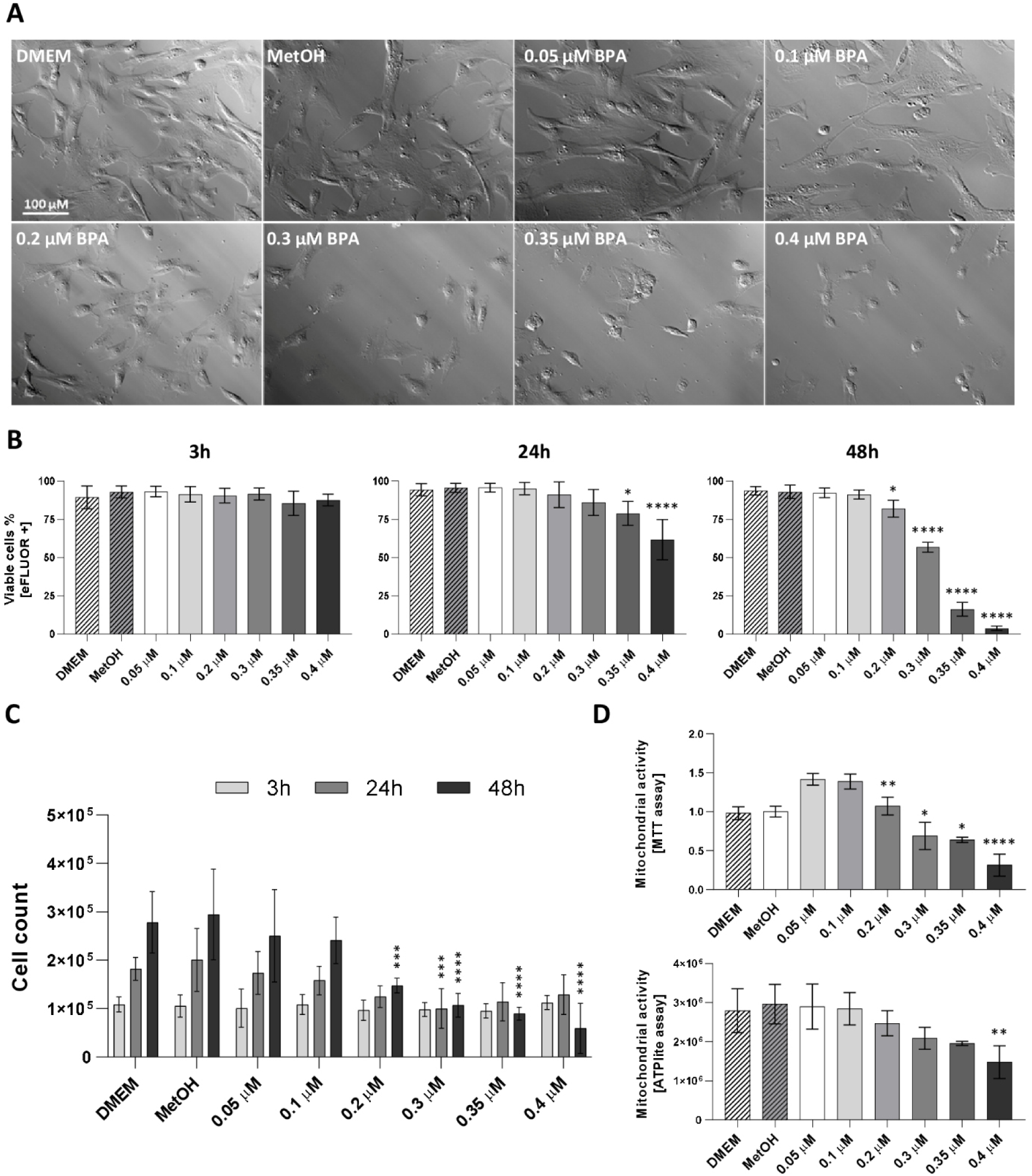
Assessment of hAMSC viability after 3, 24 and 48 hours of exposure to increasing BPA concentrations. Morphological differences in hAMSC after 24 hours of exposure with increasing concentrations of BPA (0.05, 0.1, 0.2, 0.3, 0.35, and 0.4 μM) (A). Viable cell quantification after 3, 24, and 48 hours of BPA exposure, conducted through eFLUOR-positive staining (B), with further validation via total cell counts obtained through flow cytometry (C). Evaluation of hAMSC vitality after 24 hours of exposure using the MTT and ATPlite assays to measure alterations in mitochondrial functionality (D). Results are represented as histograms showing mean values ± SD from n≥3 independent experiments. Statistical analysis was performed versus the control condition represented by MetOH: p < 0.01(**\*\***), p < 0.001(**\*\*\****), p < 0.0001(*\*\*\*\*)).

The impact of increasing concentrations of BPA on hAMSC viability was investigated 3, 24, and 48 hours after exposure to increasing concentrations of BPA by quantifying the percentage of eFluor-positive cells. While no significant differences were observed after 3 hours (Figure 1B, left panel), a noticeable effect on cellular viability was observed after 24 hours, starting from 0.35 μM BPA treatment (Figure 1B, central panel). This effect intensified at 48 hours, with significant reductions starting from 0.2 μM BPA treatment (Figure 1B, right panel). These findings were corroborated by flow cytometry’s absolute cell count measurements. As shown in Figure 1C, a marked reduction in total cell count began after 24 hours of exposure, reaching statistical significance at 0.3 μM BPA concentration. This trend persisted 48 hours after exposure, with significant results starting from 0.2 μM BPA concentration. No variations were observed at 3 hours (Figure 1C).

Furthermore, we assessed the impact of BPA on cellular viability using the MTT and ATPlite assays, which evaluate mitochondrial activity and function. Results from both assays demonstrated a pronounced reduction in cellular viability, with significance observed from 0.3 μM BPA treatment in the MTT, and only at the highest tested concentration (0.4 μM) in the ATPlite assay.

### 2. Increasing BPA concentrations induce oxidative stress in hAMSC initiating an antioxidant response

Results obtained by the MTT and ATPlite assays indicated changes in mitochondrial activity. Therefore, we assessed whether increasing concentrations of BPA affected mitochondrial production of ROS (mtROS). The evaluation was conducted 3 and 24 hours after BPA exposure, excluding the 48-hour time point due to pronounced toxicity at the highest tested concentrations. mtROS was quantified using the Mitosox Red fluorescent dye in flow cytometry and immunofluorescence.

Three hours after exposure, flow cytometry analysis revealed a slight increase in Mitosox Red-positive cells, with significant values reached starting from 0.35 μM BPA concentrations (Figure 2A, left panel). These results were more pronounced after 24 hours (Figure 2A, right panel).

**Figure 2.**
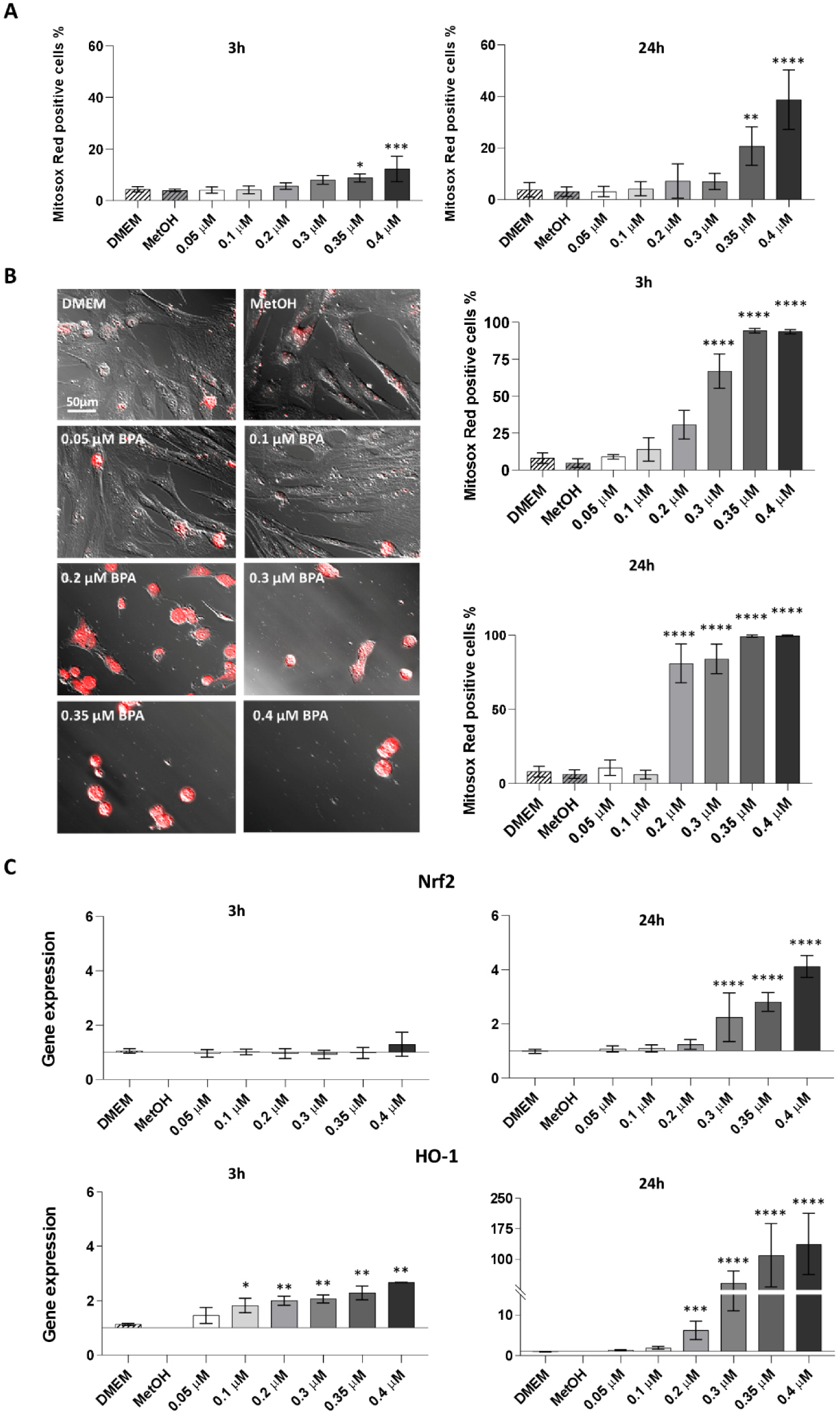
Evaluation of hAMSC oxidative stress after 3 and 24 hours of exposure to increasing BPA concentrations. Mitochondrial reactive oxygen species (mtROS) production in hAMSC was quantified in flow cytometry and in immunofluorescence using the Mitosox Red fluorescent dye, after 3 and 24 hours of exposure to increasing concentrations of BPA (0.05, 0.1, 0.2, 0.3, 0.35, and 0.4 μM). Results acquired in flow cytometry are presented as the percentage of Mitosox Red-positive cells (Figure 2A). Immunofluorescent images were acquired at 20x magnification and Mitosox Red-positive cells were identified with red signal (Figure 2B, 50 μm scale bar). The antioxidant response of hAMSC was assessed by measuring Nrf2 and HO-1 gene expression after 3 and 24 hours of BPA exposure, expressed as fold-change relative to the control condition (MetOH) (Figure 2C). Histograms represent mean values ± SD from n=4 independent experiments. Statistical analysis was performed versus the control condition: p < 0.01(**), p < 0.001(***), p < 0.0001(****).

Similarly, immunofluorescence images showed more red-fluorescent cells after 3 and 24 hours of exposure to increasing BPA concentrations (Figure 2B). There were no changes observed at both time points for MetOH-treated, DMEM-cultured, and cells exposed to lower BPA concentrations (0.05 and 0.1 μM BPA). However, Mitosox Red positivity became significant after exposure to 0.3 μM BPA at 3 hours (Figure 2B, upper panel; immunofluorescence images are in Supplementary Figure 1) and 0.2 μM BPA at 24 hours (Figure 2B, bottom panel).

In response to oxidative stress, cells attempt to counteract ROS by implementing an antioxidant response. Here we assessed the expression of Nrf2, a critical transcription factor involved in protecting against oxidants, and its principal effector protein, HO-1 (27), (28). Three hours after exposure, we observed a mild variation in Nrf2 gene expression at the highest BPA concentration tested, alongside a significant upregulation of HO-1 gene expression (Figure 2C, left panels). This effect was more pronounced after 24 hours, with Nrf2 gene upregulation starting from 0.3 μM BPA, and a significant HO-1 expression from 0.35 μM BPA (Figure 2C, right panels).

Furthermore, as no substantial differences were observed between the DMEM and MetOH control condition, subsequent experiments were performed only with the MetOH condition.

### 3. Increasing concentrations of BPA modulate the expression of inflammasome-related genes in hAMSC, without affecting IL-1**β** secretion

Mitochondrial reactive oxygen species (mtROS) is a key activator of the inflammasome (29). Considering the significant increase in mtROS production (Figure 2A and 2B), we investigated the recruitment and the activation of the inflammasome complex in hAMSC exposed to increasing BPA concentrations. Particularly, we assessed the genetic expression of structural and effector components from 3 inflammasomes (NLRP3, NLRC4, AIM2) after 24 hours of BPA exposure. Among these sensor proteins, which form multimeric complexes in response to stimuli in the cytoplasm, NLRP3 gene expression significantly increased with 0.2 and 0.3 μM BPA concentrations, returning to control levels at higher BPA concentrations (Figure 3). Differently, genes encoding for NLRC4 and AIM2 sensor proteins, showed a significant upregulation following treatment with higher BPA concentrations (0.3; 0.35 and 0.4 μM). These results suggest varied effects of BPA on the diverse inflammasome complexes. Similar trends were observed for two other key players of the inflammasome, ASC and caspase 1, which exhibited significant upregulation with exposure to 0.35 μM and 0.2 μM BPA, respectively (Figure 3). IL-1β, the final effector of the inflammasome cascade, showed a genetic expression pattern similar to NLRP3, with a significant increase compared to control after exposure to 0.2 μM BPA, and a significant decrease after exposure to 0.35 and 0.4 μM BPA (Figure 3). However, unlike the genetic findings, IL-1β protein expression was undetected after exposure to increasing BPA concentrations, as assessed by ELISA (Supplementary Figure 2). The lack of IL-1β production may indicate that BPA acts solely as a priming agent in inflammasome complex activation, where gene transcription occurs. Nevertheless, even after exposing the cells to a second stimulus adenosine triphosphate (ATP) to complete the activation of the complex (30),(31), IL-1β production was not observed (Supplementary Figure 2).

**Figure 3.**
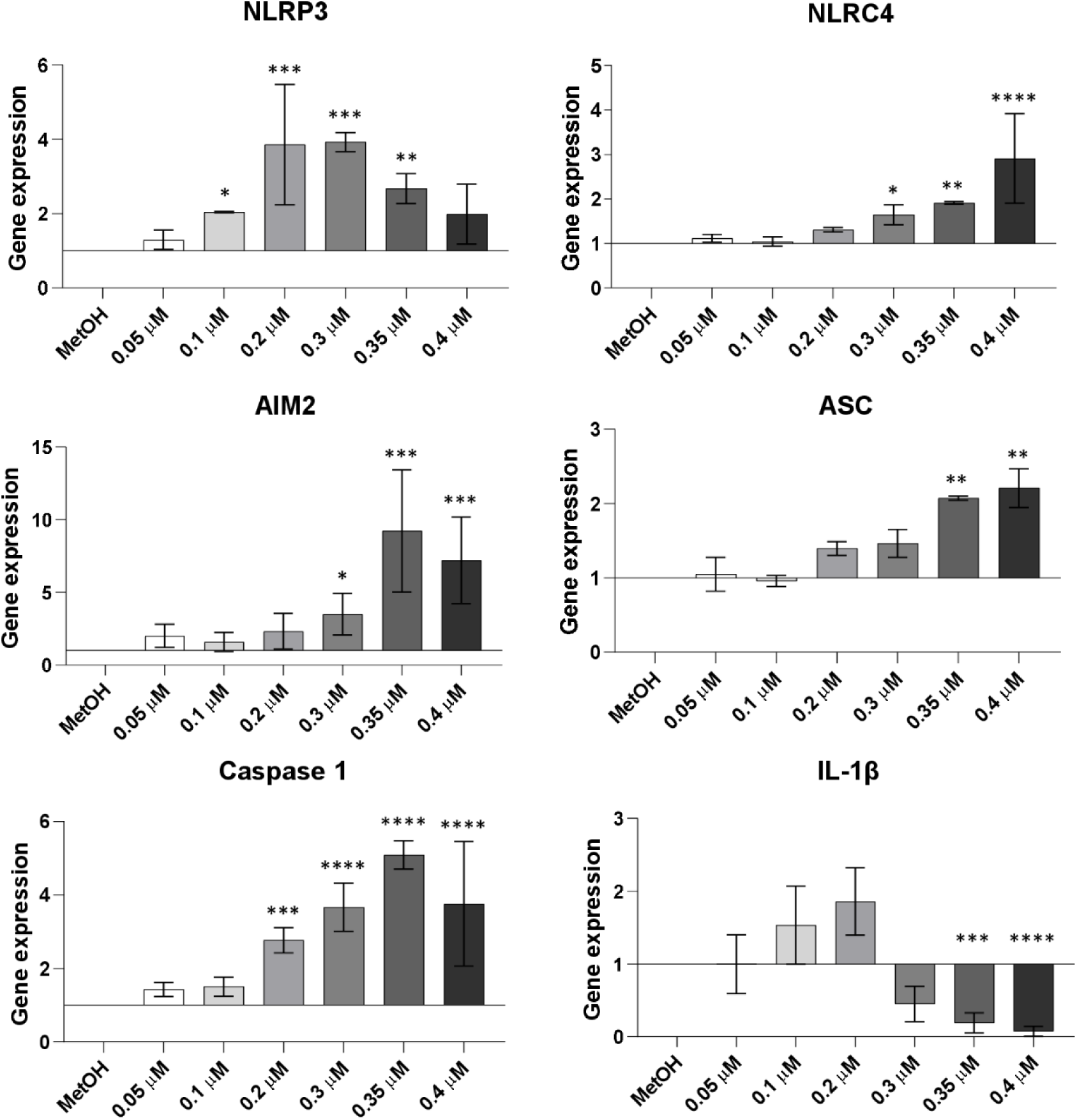
Evaluation of inflammasome recruitment and activation in hAMSC after 24 hours of exposure to increasing BPA concentrations. Inflammasome formation and activation were assessed by examining the genetic expression of both structural and functional components of the complex (NLRP3, NLRC4, AIM2, ASC, caspase 1, IL-1β) (Figure 3) and the secretion of effector protein (IL-1β) (Supplementary) after exposure to increasing BPA concentrations (0.05, 0.1, 0.2, 0.3, 0.35, and 0.4 μM) with or without ATP (1.5 mM). The figure illustrates alterations in gene expression, presented as fold-change relative to the control condition (MetOH). Histograms represent the mean values ± SD from n=3 independent experiments. Statistical analysis was performed versus the control condition: p < 0.01(**), p < 0.001(***), p < 0.0001(****).

### 4. Increasing concentrations of BPA induce the gene expression of p53, p21 and p27 genes in hAMSC

In response to oxidative stress, cells can activate specific damage-response mechanisms, leading to cell cycle arrest, and potentially inducing senescence (32). To investigate whether increased ROS production triggers these defensive mechanisms in hAMSC, we examined the expression of p53 gene, a primary transcription factor recruited under stressful conditions. Results from RT-PCR revealed a significant upregulation in p53-encoding gene, starting from 24 hours of exposure with 0.3 μM BPA (Figure 4A, left panel). Furthermore, we investigated changes in the expression profiles of p21 and p27, two master transcriptional factors responsible for cell cycle arrest and senescence induction. Indeed, the recruitment of these two cycline-dependent kinase (CDK) inhibitors is often mediated by p53 expression (33). As depicted in Figure 4A (central and right panels), significant upregulation of p21 and p27 genes was observed, with significant values reached from 0.3 μM BPA concentration for p21, and only at the highest tested concentration for p27. To determine whether the increased p21 gene expression correlated with functional activity, we assessed its nuclear translocation by immunofluorescence analysis after exposure to increasing BPA concentrations (0.1, 0.2, 0.3, 0.4 μM). As observed in Figure 4B, increased nuclear translocation of p21 was evident even at the lowest concentration tested, with significance reached only at the highest tested concentration of 0.4 μM BPA.

**Figure 4.**
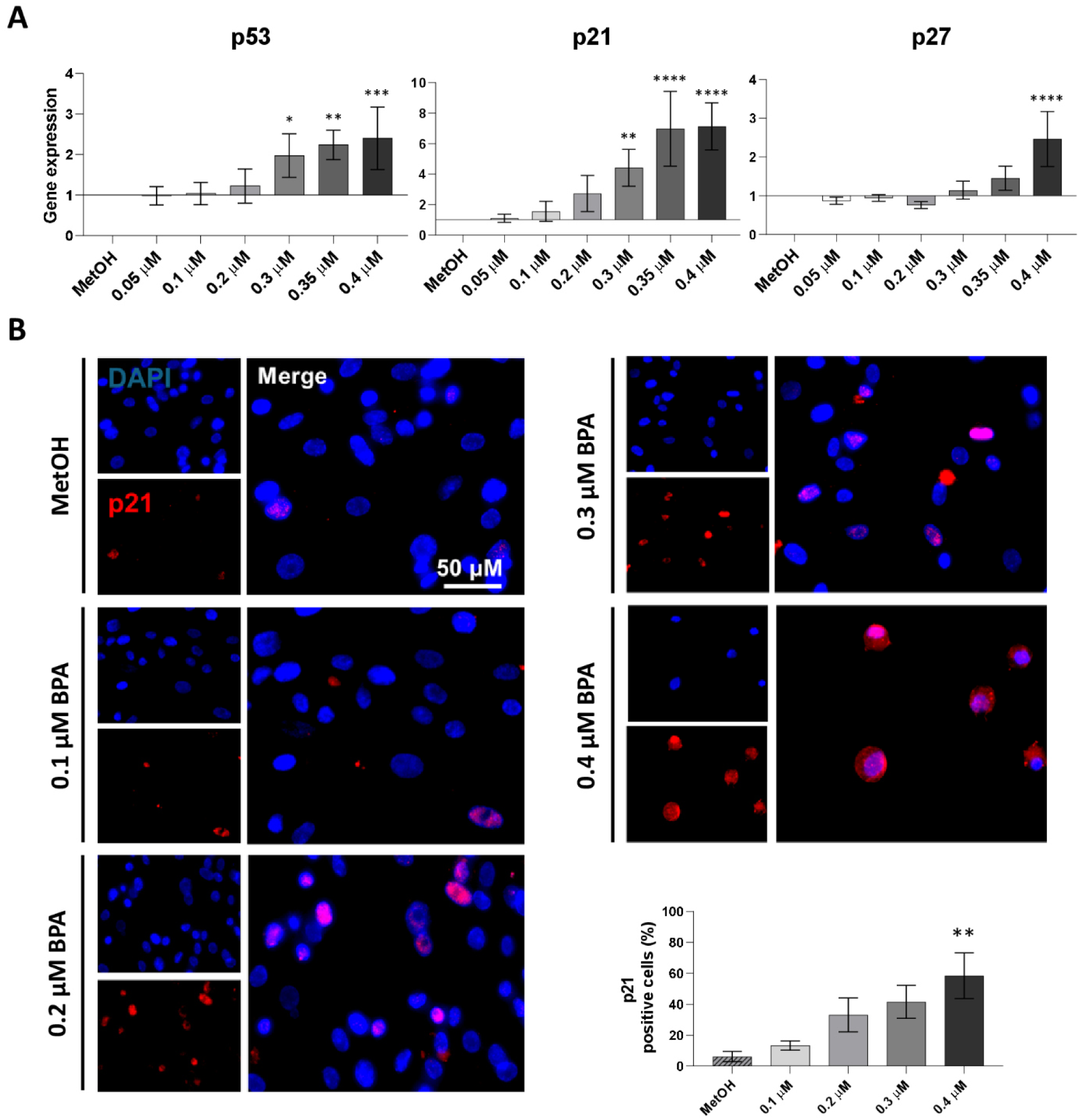
Investigation of cell cycle gene expression in hAMSC after 24 hours of exposure to increasing BPA concentrations. Expression of p53, p21 and p27 cell-cycle regulating genes was analysed by RT-PCR 24 hours after exposure to increasing BPA concentrations (0.05, 0.1, 0.2, 0.3, 0.35 and 0.4 μM). Results are presented as fold-change relative to control conditions (MetOH) (Figure 4A). Furthermore, p21 protein expression and its nuclear translocation in hAMSC were assessed 24 hours after exposure to increasing BPA concentrations (0.1, 0.2, 0.3, and 0.4 μM) using immunofluorescence analysis (Figure 4B). p21-positive cells were identified by a rosy-red signal, while nuclei were stained with DAPI (blue). Pictures were acquired at 20x magnification (scale bar corresponds to 50 μm in panel C). Histograms represent the mean values ± standard deviation from n=4 (Figure 4A) and n=3 (Figure 4B) independent experiments. Statistical analysis was performed versus the control condition: p < 0.01(**), p < 0.001(***), p < 0.0001(****).

### 5. Increasing concentrations of BPA induce senescence in hAMSC with the acquisition of the SASP profile

To validate our findings and confirm the acquisition of a senescent phenotype in hAMSC, we assessed β-galactosidase activity in cells treated with increasing BPA concentrations. Results were compared with those obtained from hAMSC treated with the epigenetic senescence inducers 5-aza-2’-deoxycytidine (5-aza) and SAHA, serving as positive controls (34). β-galactosidase (β-gal)-positive cells exhibited green perinuclear staining, with a higher frequency and intensity in cells exposed to 0.3 and 0.4 μM BPA concentrations compared to the negative control (MetOH-treated cells), and similar to the positive controls. As shown in the representative histograms (Figure 5A), the percentage of senescent cells reached significance starting from 0.2 μM BPA concentration and exceeded 70% positivity for hAMSC exposed to 0.4 μM BPA concentration (as observed in 5-aza and SAHA-treated cells).

**Figure 5.**
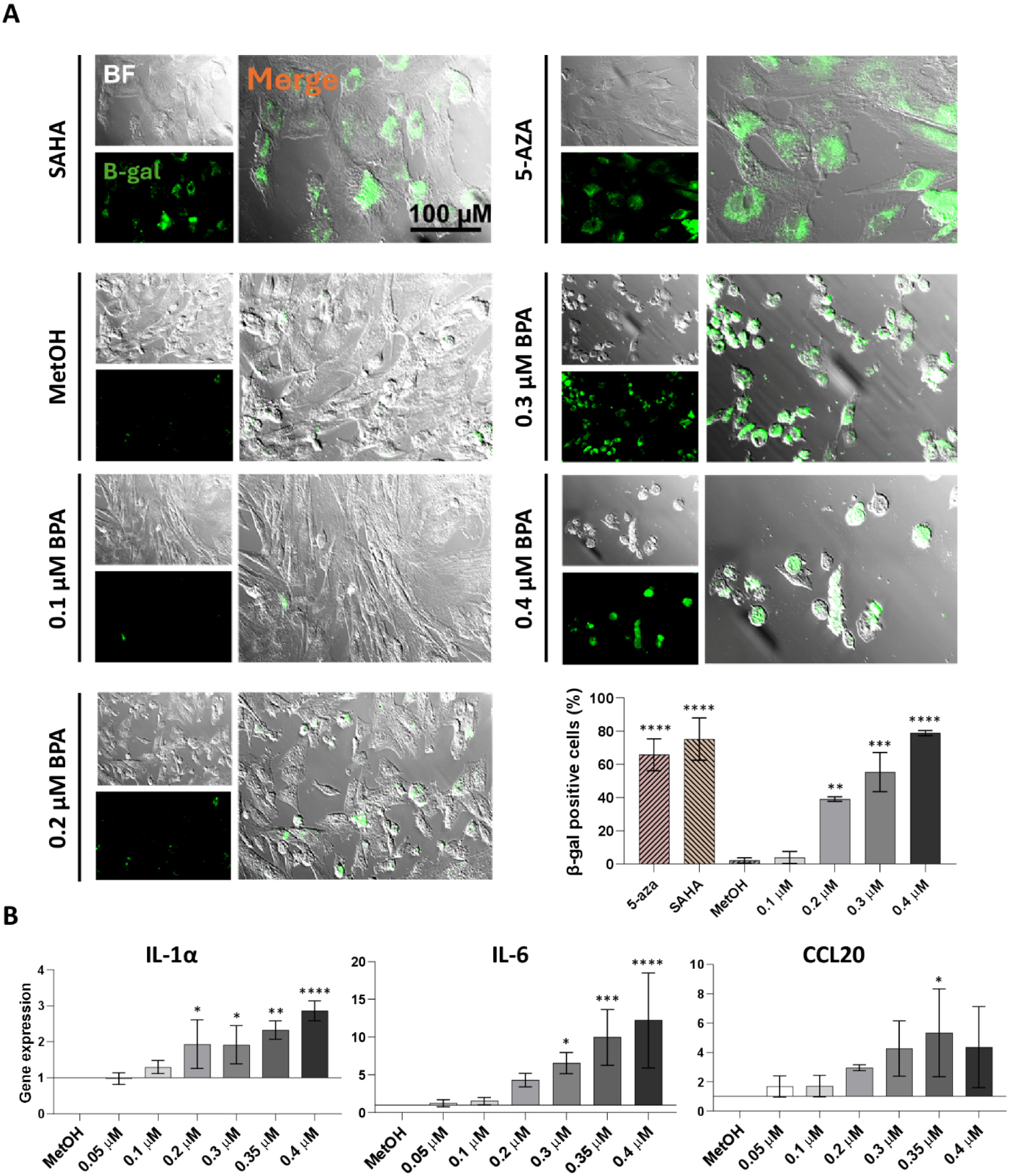
Investigation of senescent induction in hAMSC after 24 hours of exposure to increasing BPA concentrations. Beta-galactosidase (β-gal) activity in hAMSC was evaluated by immunofluorescence analysis 24 hours after exposure to increasing concentrations of BPA (0.1, 0.2, 0.3, and 0.4 μM) (Figure 5A). β-gal-positive cells were identified by a green signal. Pictures were acquired at 20× magnification (scale bar corresponds to 100 μM). Expression of senescent-associated secretory phenotype (SASP) molecules (IL-6, IL-1α and CCL20) were analysed by RT-PCR 24 hours after BPA exposure and presented as fold-change relative to the control (MetOH) (Figure 5B). Results are represented as histograms showing mean values ± SD from n=3 independent experiments. Statistical analysis was performed versus the control condition: p < 0.01(**), p < 0.001(***), p < 0.0001(****).

Since the senescence state is characterized by the secretion of bioactive factors, collectively known as senescence-associated secretory phenotype (SASP) (35), we further analysed the expression of IL-6, IL-1α, and CCL20. Similarly, to the acquisition of β-gal positivity (Figure 5A), a significant upregulation in the genetic expression was observed starting from the concentration of 0.2 μM BPA for IL-1α (Figure 5B, left panel), 0.3 μM for IL-6 (Figure 5B, central panel), and 0.35 μM for the chemokine CCL20 (Figure 5B, right panel).

### 6. Toxicity induced by increasing concentrations of BPA culminates in the induction of apoptosis with dysregulated expression of Bax and Bcl-2 genes in hAMSC

Oxidative stress can lead to cell death when all activated processes are insufficient to repair the damage (36). In this regard, we assessed the extent of cells undergoing activation of the apoptotic pathway after exposure to increasing concentrations of BPA, using the PI-Annexin V assay. Analyses conducted 3, 24, and 48 hours after exposure to BPA revealed no variations at 3 hours (Figure 6A, left panel). However, 24 hours after BPA exposure, the percentage of live cells began to decline, reaching significance starting from exposure with 0.3 μM BPA concentration. Concurrently, the number of late apoptotic cells significantly increased at the same BPA concentration (Figure 6A, central panel). This effect intensified at 48 hours, with a significant reduction in the percentage of live cells observed from 0.2 μM BPA and a significant increase in late apoptotic cells from 0.3 μM BPA (Figure 6A, right panel). No changes in the percentage of early apoptotic cells were detected 3, 24, and 48 hours after BPA treatment.

**Figure 6.**
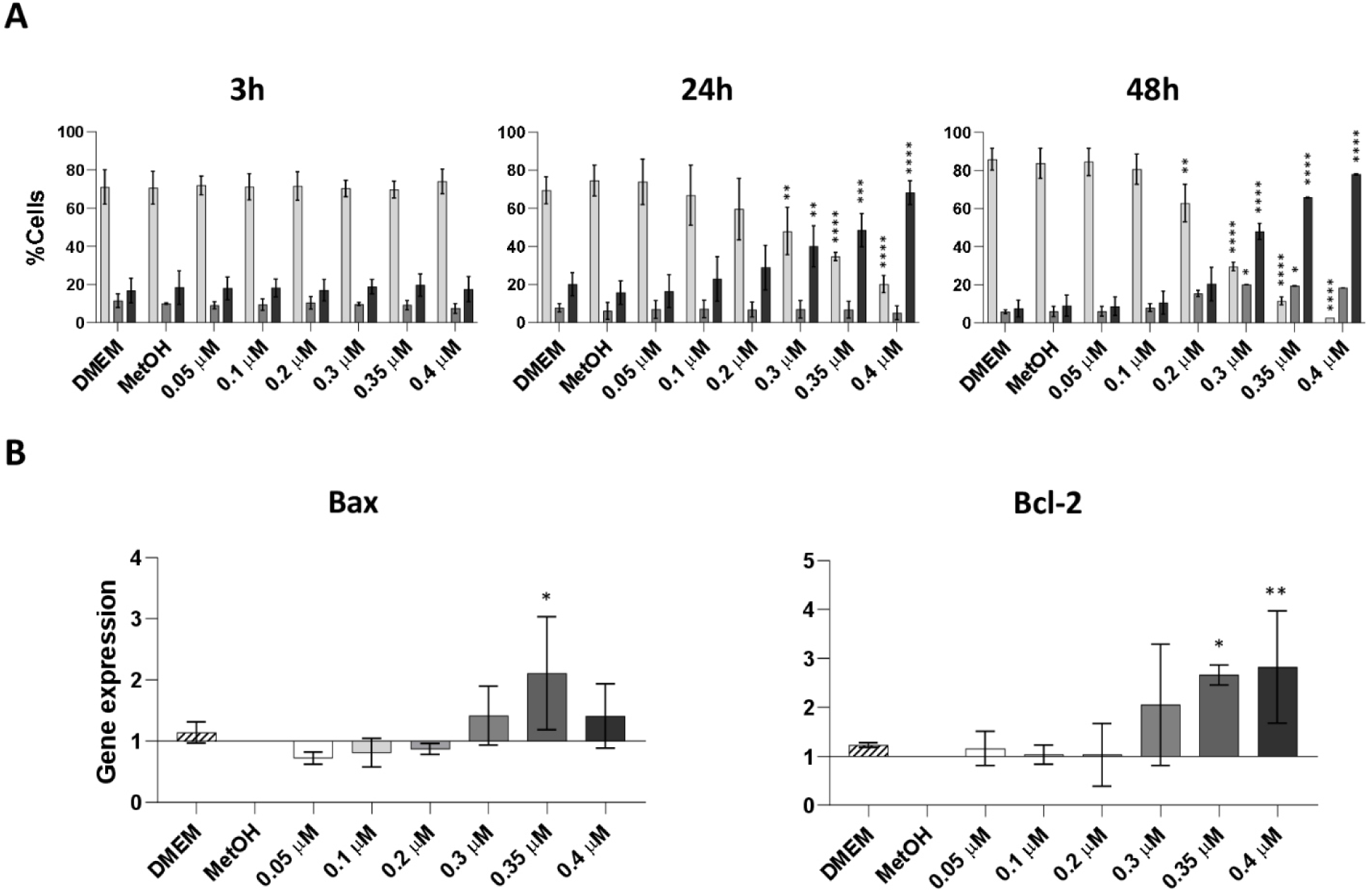
Investigation of apoptosis induction in hAMSC after 3, 24 and 48 hours of exposure to increasing BPA concentrations. Percentage of live, early and late apoptotic cells was measured by Annexin-PI fluorogenic dyes 3, 24 and 48 hours after exposure to increasing concentrations of BPA (0.05, 0.1, 0.2, 0.3, 0.35, and 0.4 μM) (Figure 6A). Expression of pro-apoptotic (Bax) and anti-apoptotic (Bcl-2) genes was analysed by RT-PCR 24 hours after BPA exposure and presented as fold-change relative to the control condition (MetOH) (Figure 6B). Results are represented as histograms showing mean values ± SD from n=3 independent experiments. Statistical analysis was performed versus the control condition: p < 0.01(**), p < 0.001(***), p < 0.0001(****).

Additionally, the genetic expression of the two well-known pro-apoptotic and anti-apoptotic genes, Bax and Bcl-2, was dysregulated after 24 hours of BPA exposure, with significant values reached with 0.35 and 0.4 μM BPA concentrations (Figure 6B). A summary diagram illustrating the impact of BPA on hAMSC is shown in Figure 7.

**Figure 7.**
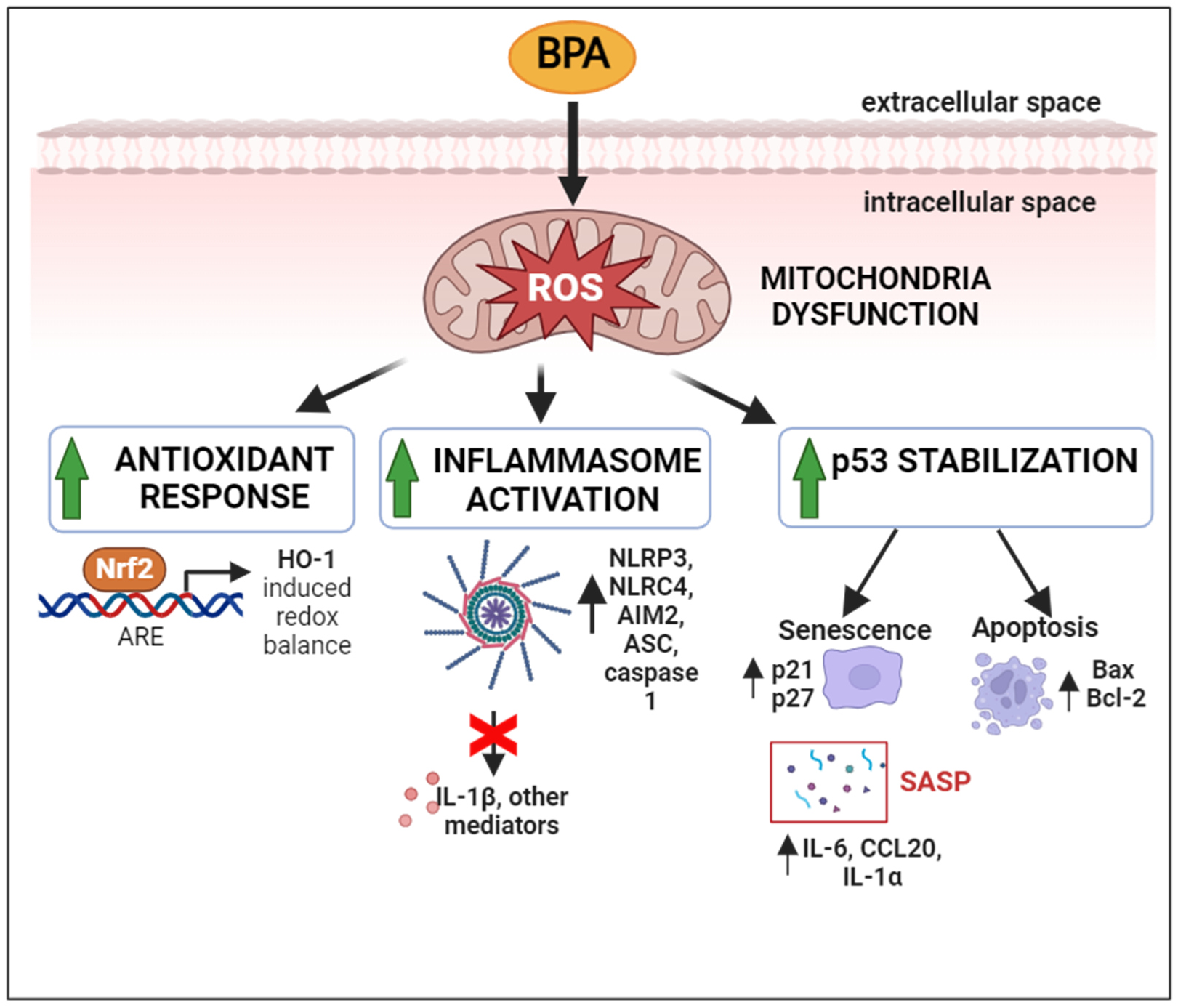
Proposed outline of the signalling pathways activated in hAMSC after exposure to increasing BPA concentrations. Alteration in mitochondrial function, evidenced by enhanced production of reactive oxygen species (ROS) in hAMSC after BPA exposure, trigger the downstream activation of several signalling pathways. ROS accumulation activates an antioxidant response, which is marked by an elevated production of Nrf2 and HO-1. Concurrently, the high ROS level induce sterile inflammation, resulting in increased transcription of factors involved in inflammasome complex formation and activation. However, this increased transcription does not translate to higher production of IL-1β, the downstream effector of the inflammasome pathway. Instead, oxidative stress promotes p53 stabilization and upregulates p21 and p27 genes, as well as components of the senescence-associated secretory phenotype (SASP). Ultimately, the senescent state serves as a prelude to apoptosis, which occurs when hAMSC are exposed to the highest BPA concentrations.

## Discussion

In recent years, the threat of microplastics to human health has been widely discussed, particularly concerning pregnant women and infants, who are especially sensitive to toxicants exposure (5). A landmark discovery in this topic was published in 2021 by Ragusa and colleagues, which demonstrates the presence of microplastics in different areas of the human term placenta (the fetal portion, the maternal portion, and the amniotic membrane) (11). Considering the crucial role of the placenta as the interface between the fetus and the external environment, the discovery of MPs within it raises concerns about the potential negative effects of microplastic pollution on pregnancy progression. In particular, MPs toxicity is mostly attributed to plasticizers (37), among which bisphenol A (BPA) has been extensively studied for its possible link with obstetric complications, such as preeclampsia, fetal growth restriction, miscarriage, and preterm birth (17). The mechanisms by which BPA influences these diseases are still unclear, prompting numerous studies to explore its intracellular actions. Our research is the first to examine the effects of BPA on mesenchymal stromal cells from the amniotic membrane (hAMSC). The aim is not only to assess BPA’s impact on these cells but also to uncover the pathways through which it operates.

We started investigating the impact of BPA on hAMSC viability, noting a significant reduction after 24 hours of exposure with the two highest BPA concentrations tested (0.35, 0.4 µM). BPA-induced viability reduction is extensively reported. Similar decreases were observed in RAW 264.7 macrophages and MLO-Y4 osteocytes exposed to 50 µM and 200 µM BPA, respectively, for 24 hours (38). Similarly, 10 µM BPA induced cytotoxicity and reduced viability in ovarian cancer cell lines after 48 hours of exposure (39), while 100 mM BPA negatively affected BeWo trophoblast viability after 24 hours of exposure (40). Although these concentrations exceed those used in our experiments, they confirm BPA’s capacity to decrease cellular viability. It noteworthy that these studies employ cell lines, which may be more resistant than primary cells like hAMSC. Notably, research on human fetal lung fibroblasts showed that BPA affected the expression of genes involved in cell cycle regulation at concentrations comparable to those used in our study (41).

The range of BPA concentrations tested in our research was chosen to closely replicate levels encountered by mothers during gestation. This selection was based on recent studies assessing serum and urine BPA concentrations in pregnant women to evaluate possible association with gestational complications (13), (12). We selected six different doses, including at least one with negligible effects (0.05 µM) and one causing a significant reduction (0.4 µM) on cell viability, comparable to the concentrations found in the blood and urine of woman with complicated pregnancies (13), (12). Furthermore, our range aligns with the *lowest observable adverse effect level* (LOAEL) defined for the in vitro evaluation of BPA toxicity, which corresponds to 2.19 x 10^^-7^ M (26), (42). In this study, we define concentrations closer to this value, ranging from nanomolar to millimolar, as *low doses*.

Our research focused on assessing whether BPA exposure induces oxidative stress in hAMSC. As expected, both early (3 hours after treatment) and late (24 hours after treatment) evaluations showed a significant increase in mtROS production at the two highest BPA concentrations tested (0.35 and 0.4 µM). BPA toxicity is extensively linked to oxidative stress, with evidence reported in vitro (43), in vivo (44), and in humans (45), (46). Similar to our findings, oxidative stress and mitochondrial dysfunction have been observed in various cell types exposed to low doses of BPA. HepG2 hepatoma cells exposed to nanomolar BPA concentrations showed increased ROS production after 24 hours (47) and reduced mitochondrial membrane potential after 6 hours (48). GT1-7 hypothalamic neurons exposed to micromolar BPA concentrations exhibited increased mtROS production after 6 hours (49), while bovine granulosa cells showed increased oxidative stress and upregulation of antioxidant enzymes when exposed to low BPA concentrations for 12 hours (43).

We also investigated dysregulation in the antioxidant response. Twenty-four hours after BPA exposure, we observed increased expression of the Nrf2 gene, along with significant upregulation of its downstream effector, the heme oxygenase-1 (HO-1). These two transcription factors are involved in cellular detoxification mechanisms (50), and their increased expression suggests an attempt to respond to BPA toxicity. The role of Nrf2 in BPA-mediated toxicity is well-documented. Human B cells exposed to 100 µM BPA underwent autophagy via Nrf2-mediated expression of pro-survival Atg7 and Beclin1 (51) and showed Nrf2 upregulation when exposed to 50 and 100 µM BPA concentrations (52). In vivo, exposure to low doses of BPA in frog embryos led to teratogenesis, DNA damage, and abnormal expression of antioxidant enzymes (SOD1, NQO1, and CAT), likely mediated by a disrupted Nrf2 signaling pathways (53). Nrf2 is crucial for human embryonic stem cell differentiation (54) and stem cell property maintenance (55). Thus, Nrf2 dysregulation may be a cellular response to mitigate BPA toxicity but could have deleterious effects on placental-resident stem cells during gestation.

As previously mentioned, heightened oxidative stress activates numerous signaling pathways, including antioxidant response, inflammation, apoptosis, and senescence ((56)). BPA-induced oxidative stress has been linked to inflammation, apoptosis, and mitochondrial dysfunction in the colon and liver of mice (57). Picomolar concentrations of BPA disrupt proliferative and inflammatory pathways in human endothelial cells through oxidative stress (58). Considering that ROS production can activate the inflammasome complex (29), we investigated whether BPA exposure induces the activation of a well-characterized inflammasome, the nucleotide-binding oligomerization domain (NOD)-like receptor containing pyrin domain 3 (NLRP3), in hAMSC. We observed upregulation of NLRP3, caspase 1, ASC, and IL-1β genes, but not the protein expression of the final effector IL-1β in hAMSC supernatant. This result suggests a blockade in the activation of the inflammasome complex. However, the complete inflammasome activation requires an initial priming stimulus which induces the expression of inflammasome-related genes, followed by a second stimulus to trigger the release of the final effectors (30), (30). In our experiments, even the addition of a second stimulus (ATP) failed to induce IL-1β production and NLRP3 inflammasome activation.

Solely two other studies have documented the involvement of NLRP3 inflammasome, in osteocytes exposed to 100 and 200 µM BPA (38) and in RAW 264.7 murine macrophages exposed to micromolar BPA concentrations (59). They show contrasting results: the former observed increased expression of inflammasome-associated proteins and induction of pyroptosis, the latter reported inactivation of the NLRP3 inflammasome and lack of IL-1β and IL-18 production. These observations highlight a different effect of BPA on this specific intracellular pathway, which nonetheless result in dysregulated cellular response in both cases. Based on our results, our hypothesis was that BPA-mediated toxicity may be severe enough to elicit a distinct cellular response.

Following exposure to the highest BPA concentrations (0.3, 0.35, and 0.4 µM), we observed upregulation of p53 gene, along with p21 and p27 cell-cycle regulators, markers typically associated with cellular senescence. p53, which plays a crucial role in determining cell fate under stress condition, can induce p21 expression, leading to senescence or apoptosis. Senescence is characterized by irreversible cell-cycle arrest and secretion of inflammatory molecules, which trigger a senescent phenotype in neighboring cells as well (35).

Herein, immunofluorescence analysis revealed nuclear localization of p21 starting from 0.2 µM BPA, the same concentration at which hAMSC showed beta-galactosidase activity. Additionally, hAMSC exhibited elevated levels of inflammatory molecules (IL-1alpha, IL-6, CCL20), typically associated with the senescence-associated secretory phenotype (SASP). These findings collectively demonstrate that increasing BPA concentrations trigger senescence in hAMSC. Similar findings were observed in primary and prostate cancer cells exposed to 10, 50, 100 µM BPA concentrations for 24, 48, and 72 hours, which showed increased p21 and p27 expression, leading to cell cycle arrest via the EGFR/ERK/p53 signaling pathway (60). Murine aortic endothelial cells exposed to 100 nm to 5 µM BPA exhibited increased beta-galactosidase activity and expression of p16 and p21, hallmarks of senescence (61). Other studies also demonstrated a positive association between BPA exposure, oxidative stress, and senescence (41), (62). In vivo experiments reported that low doses of BPA induce renal injury in adult rats, evidenced by histological alterations and increased cellular senescence (63).

In the context of BPA exposure, the induction of senescence state in hAMSC can be considered an adaptive stress response. Moreover, it is important to highlight that senescence is a required process during embryological development, but excessive senescence can be associated with negative pregnancy outcomes (64).

hAMSC may respond to BPA-induced oxidative stress by arresting cell cycle progression and proliferation, as evidenced by the dramatic decrease in cell numbers at higher BPA concentrations.

As suggested by the PI/Annexin evaluation, BPA-mediated toxicity can activate apoptosis starting from 24 hours of exposure. The upregulation of pro-apoptotic Bax and anti-apoptotic Bcl-2 genes in response to the higher BPA concentrations (0.3, 0.35, and 0.4 µM), suggests their involvement in apoptosis induction. Interestingly, our findings did not indicate a necrotic process, as suggested by PI/Annexin staining, and the release of inflammatory molecules appears to be linked solely to the senescent state.

In conclusion, our results, consistent with current research, suggest that BPA is an environmental pollutant extremely toxic to cells, and specifically also on MSC derived from amniotic membrane (hAMSC). Our findings go beyond current research by demonstrating, for the first time, the in vitro impact of environmentally relevant concentrations of BPA on hAMSC, elucidating pathways through which it acts. We observed a reduction in hAMSC viability and increased production of mtROS, associated with mitochondrial dysfunction. The increased oxidative stress appears to play a central role, triggering a series of downstream signaling pathways aimed to restore cellular homeostasis. Additionally, our in vitro evaluations suggest that hAMSC exposed to increasing BPA concentrations, partially arrest cell cycle progression, entering senescence, and partly undergo apoptosis, when toxicity is unsustainable. These results contribute to a broader understanding of BPA’s direct impact on placenta cells.

Considering the limitations inherent in investigations during pregnancy, these findings are significant for informing large-scale decisions, such as restricting BPA in certain products to safeguard the health of both the mother and her baby, as well as public health overall.

## Acknowledgments

The authors would like to thank the Regenerative Medicine Research Center (CROME) of Università Cattolica del Sacro Cuore, the physicians and midwives of the Department of Obstetrics and Gynaecology of Fondazione Poliambulanza-Istituto Ospedaliero, Brescia (Italy), and all the mothers who donate their baby’s placenta to make this research possible.

## Author contribution

Conceptualization A.P. and O.P.; methodology S.F, A.P., M.M.; investigation S.F., A.P., M.M.; writing – original draft S.F., A.P., A.S., and O.P.; writing – review & editing S.F., A.P., A.M., M.M, M.G., A.S., and O.P.; funding acquisition O.P.; supervision A.P. and O.P.

